# 3D U-Net improves automatic brain extraction for isotropic rat brain MRI data

**DOI:** 10.1101/2021.09.27.462020

**Authors:** Li-Ming Hsu, Shuai Wang, Lindsay Walton, Tzu-Wen Winnie Wang, Sung-Ho Lee, Yen-Yu Ian Shih

**Author notes:** Equal contribution (Li-Ming Hsu, Shuai Wang). **Correspondent author**: Li-Ming Hsu, Yen-Yu Ian Shih.

## Abstract

Brain extraction is a critical pre-processing step in brain magnetic resonance imaging (MRI) analytical pipelines. In rodents, this is often achieved by manually editing brain masks slice-by-slice, a time-consuming task where workloads increase with higher spatial resolution datasets. We recently demonstrated successful automatic brain extraction via a deep-learning-based framework, U-Net, using 2D convolutions. However, such an approach cannot make use of the rich 3D spatial-context information from volumetric MRI data. In this study, we advanced our previously proposed U-Net architecture by replacing all 2D operations with their 3D counterparts and created a 3D U-Net framework. We trained and validated our model using a recently released CAMRI rat brain database acquired at isotropic spatial resolution, including T2-weighted turbo-spin-echo structural MRI and T2*-weighted echo-planar-imaging functional MRI. The performance of our 3D U-Net model was compared with existing rodent brain extraction tools, including Rapid Automatic Tissue Segmentation (RATS), Pulse-Coupled Neural Network (PCNN), SHape descriptor selected External Regions after Morphologically filtering (SHERM), and our previously proposed 2D U-Net model. 3D U-Net demonstrated superior performance in Dice, Jaccard, Hausdorff distance, and sensitivity. Additionally, we demonstrated the reliability of 3D U-Net under various noise levels, evaluated the optimal training sample sizes, and disseminated all source codes publicly, with a hope that this approach will benefit rodent MRI research community.

**Significant methodological contribution:** We proposed a deep-learning-based framework to automatically identify the rodent brain boundaries in MRI. With a fully 3D convolutional network model, 3D U-Net, our proposed method demonstrated improved performance compared to current automatic brain extraction methods, as shown in several qualitative metrics (Dice, Jaccard, PPV, SEN, and Hausdorff). We trust that this tool will avoid human bias and streamline pre-processing steps during 3D high resolution rodent brain MRI data analysis. The software developed herein has been disseminated freely to the community.

## Introduction

Magnetic resonance imaging (MRI) is a commonly utilized technique to noninvasively study the anatomy and function of rodent brains (Mandino et al., 2019). Among the data pre-processing procedures, brain extraction is an important step that ensures the success of subsequent registration processes (Uhlich et al., 2018). Automating brain extraction is particularly challenging for rodent brains as compared to humans because of differences in brain/scalp tissue geometry, image resolution with respect to brain-scalp distance, and tissue contrast around the skull (Hsu et al., 2020). Therefore, in practice, this is often achieved by manually drawing brain masks for every slice, making it a time-consuming and operator-dependent process. Datasets with high through-plane resolution make manual brain extraction a particularly daunting task. Therefore, a robust and reliable automatic brain extraction tool would streamline the pre-processing pipeline, avoid personnel bias, and significantly improve research efficiency (Babalola et al., 2009; Feo and Giove, 2019; Gaser et al., 2012; Hsu et al., 2020; Lu et al., 2010).

To date, the most prominent tools to address rodent MRI brain extraction include Pulse-Coupled Neural Network (PCNN)-based brain extraction proposed by (Chou et al., 2011), Rapid Automatic Tissue Segmentation (RATS) proposed by (Oguz et al., 2014), and SHape descriptor selected External Regions after Morphologically filtering (SHERM) proposed by (Liu et al., 2020), as well as a convolutional deep-learning based algorithm, 2D U-Net, proposed by (Hsu et al., 2020). While PCNN, RATS, and SHERM have demonstrated remarkable success (detailed introduction and comparisons discussed in (Hsu et al., 2020)), their performance is subject to brain size, shape, texture, and contrast; hence, their settings often need to be adjusted per MRI-protocol for optimal results. In contrast, the U-Net algorithm explores and learns the hierarchical features from the training dataset, and provides a user-friendly and more universally applicable platform (Ronneberger et al., 2015; Yogananda et al., 2019).

As a specific type of convolutional neural network (CNN) architecture (Krizhevsky and Sutskever, 2012), U-Net has proven valuable in biomedical image segmentation (Ronneberger et al., 2015; Yogananda et al., 2019). U-Net utilizes the encoder/decoder structure that easily integrates multi-scale information and has better gradient propagation during training. Our previous 2D U-Net approach uses 2D convolutional kernels to predict brain boundaries on a single slice basis. However, since the 2D framework only takes a single slice as input and does not utilize information across slice direction, it inherently fails to leverage context from adjacent slices in a volumetric MRI dataset. To improve upon and possibly outperform 2D U-Net, a 3D U-Net framework using 3D convolutional kernels to predict segmentation predictions on volumetric patches must be explored (Çiçek et al., 2016a).

In this work, we demonstrated the use of 3D U-Net for brain extraction in high resolution 3D rat brain MRI data. The whole network is implemented based on Keras (Chollet, 2015) with TensorFlow (Martín et al., 2016) as the backend. We trained and tested the 3D U-Net model for brain extraction performance using a recently released CAMRI rat brain database (Lee et al., 2021), including T2-weighted (T2w) rapid acquisition with relaxation enhancement (RARE) structural MRI (0.2 mm isotropic resolution) and T2*-weighted (T2*w) echo-planar-imaging (EPI) functional MRI (0.4 mm isotropic resolution). The 3D U-Net model was compared with existing rodent brain extraction tools, including RATS, PCNN, SHERM, and 2D U-Net. To benchmark the utility of this approach to rodent MRI data of various quality, we assessed 3D U-Net performance under various noise levels and evaluated the optimal training sample sizes. It is our hope that the information provided herein, together with dissemination of source codes, will benefit the rodent MRI research community.

## Methods

### Dataset Descriptions

This study utilizes a recently disseminated CAMRI dataset (Lee et al., 2021), available at https://doi.org/10.18112/openneuro.ds003646.v1.0.0. The CAMRI dataset consisted of rats aged P80-210 weighing 300-450 g across cohorts of three different rat strains: Sprague-Dawley (n = 41), Long-Evans (n = 13), and Wistar (n = 33); and both sexes: male (n = 65) and female (n = 22). Detailed information about each cohort can be found in (Lee et al., 2021). Each subject contains a T2w RARE and an T2*w EPI, and resolutions were 0.2 mm isotropic and 0.4 mm isotropic, respectively. To train our U-Net model, we first established a training dataset by randomly selecting 80% of the T2w RARE and T2*w EPI images in the CAMRI rat database (69 subjects), leaving the remaining 20% of the data as final performance testing dataset (18 subjects). In the training process, we further randomly selected 80% of the data from the training dataset (55 subjects). The remaining 20% of the data from the training dataset was used to validate the training of U-Net model (14 subjects). We repeated the training-validation process five times to avoid potential bias in data splitting. For each U-Net algorithm, the U-Net model with the highest averaged validation accuracy was then used as the final model for testing (Fig. S2).

### U-Net

The 3D U-Net framework is shown in Figure 1. The contracting path includes 32 feature maps in the first convolutional block, 64 in the second, then 96, 128, and 256 in the third, fourth, and fifth, respectively. Compared to the configuration described by (Ronneberger et al., 2015), we replaced the cross-entropy loss function with the Dice coefficient loss (Wang et al., 2020) to free the optimization process from a class-imbalance problem (Milletari et al., 2016). Since we evaluated T2w RARE and T2*w EPI data, we performed spatial normalization for distinct resolutions. For spatial normalization, we resampled all images into the same spatial resolution at 0.2 mm × 0.2 mm × 0.2 mm using nearest-neighbor (NN) interpolation. The NN was chosen because it is most suitable for binary images (brain mask). In U-Net training, the voxels belonging to the rat brain were labeled as 1 and other voxels (background) were labeled as 0. Our network was implemented using Keras (Chollet, 2015) with TensorFlow (Martín et al., 2016) as the backend. The initial learning rate and batch size were 1e^−3^ and 16, respectively. We used Adam (Kingma and Ba, 2015) as the optimizer and clipped all parameter gradients to a maximum norm of 1. During the training process, we randomly cropped 64×64×64 sized patches from all directions as the input. During the inference process, the extracted and overlapped patches were input into the trained model with a 16×16×16 stride. The overlapped predictions were averaged and then resampled back to the original resolution using nearest-neighbor interpolation to generate final output. We also trained 1) 2D U-Net with patch size of 64×64 (2D U-Net64) and a 16×16 stride (Hsu et al., 2020) and 2) 3D U-Net with patch size of 16×16×16 and a 4×4 stride (3D U-Net16) to compare the segmentation performance with the proposed 3D U-Net of patch size 64×64×64 (3D U-Net64).

**Figure 1.**
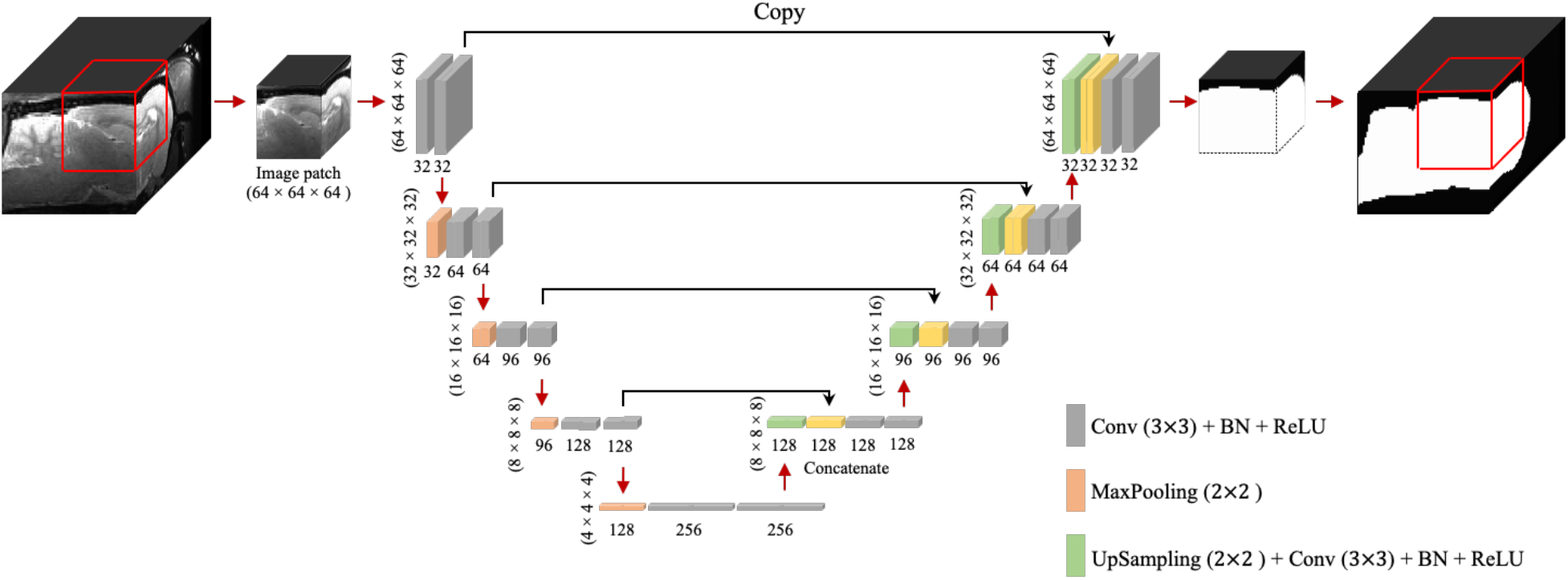
3D U-Net architecture. Boxes represent cross-sections of square feature maps. Individual map dimensions are indicated on lower left, and number of channels are indicated below the dimensions. The leftmost map is a 64 × 64 × 64 normalized MRI image patched from the original MRI map, and the rightmost represents binary ring mask prediction. Red arrows represent operations specified by the colored box, while black arrows represent copying skip connections. Conv: Convolution, BN: Batch Normalization, ReLU: Rectified Linear Unit.

### Reproducibility of 3D U-Net performance

To evaluate the 3D U-Net64 performance reproducibility, we reexamined its accuracy under two conditions: 1) Adding different Gaussian noise in the testing images, and 2) using a different number of rats from the original cohort in the training process. Specifically, as for the first validation analysis, we added Gaussian white noise in the normalized testing images with variance from 5×10^−4^ to 5×10^−5^ with step 5×10^−5^ to investigate the segmentation performance. For each testing image, the signal-to-noise ratio (SNR) was estimated to represent the image noise levels by calculating the ratio of signal intensities in the area of interest and the background (Figure S1). We estimated the average signal by using two spherical volumes of interest (VOI) with diameter = 1 mm in bilateral striatum. We also put two VOIs at background to extract noise in standard deviation of signal. To evaluate the optimal subject number in 3D U-Net64 training process, we randomly selected 5–55 training subjects in increments of 5 subjects from the total 55 training datasets, and 2, 8, and 14 validation subjects from the total 14 validation datasets. This random selection was repeated 5 times to avoid bias. We then calculated accuracy of the derived brain masks across each testing dataset.

### Evaluation Methods

To demonstrate the reliability of our proposed method, we compared our 3D U-Net method with the most prominently used methods for rat brain segmentation: RATS (Oguz et al., 2014), PCNN (Chou et al., 2011), SHERM (Liu et al., 2020), and 2D U-Net (Hsu et al., 2020). All images were bias-corrected for field inhomogeneities using Advanced Normalization Tools (N4ITK) (Avants et al., 2009; Tustison et al., 2010). The parameters in each method were chosen according to the best parameters suggested in the literature. For the RATS algorithm, the intensity threshold (T) was set to the average intensity in the entire image and the brain size value (Vt) was set to 1650 mm^3^ (Oguz et al., 2014). For the PCNN algorithm, the brain size range was set to 1000–3000 mm^3^ (Chou et al., 2011). For SHERM, the brain size range was set to 500–1900 mm^3^ (Liu et al., 2020). The convexity threshold in SHERM, defined as the ratio between the volume of a region and that of its convex hull, was set to 0.5.

To quantitatively evaluate the segmentation performance of 3D U-Net64, 2D U-Net64, 3D U-Net16, RATS, PCNN, and SHERM, we estimated the similarity of the generated brain segmentation results compared to manual brain masks (ground truth) drawn by an anatomical expert according to the Paxinos and Watson rat atlas (Paxinos and Watson, 2007). The manual segmentation was performed at the original MRI resolution before data resampling to 0.2 mm × 0.2 mm × 0.2 mm for U-Net training. Evaluations included: (1) volumetric overlap assessments via Dice, the similarity of two samples; (2) Jaccard, the similarity of two samples where Dice doesn’t satisfy the triangle inequality; (3) positive predictive value (PPV), the rate of true positives in prediction results; (4) sensitivity (SEN), the rate of true positives in manual delineation; and (5) a surface distance assessment by Hausdorff distance, the distance of two samples. The following definitions were used for each: *Dice* = 2(│*A* ∩ *B*│)/(│*A*│ + │*B*│), *jaccard* = (│*A* ∩ *B*│)/(│*A* ∪ *B*│), *PPV* = (│*A* ∩ *B*│)/*B*, *SEN* = (│*A* ∩ B│)/*A*, and *Hausdorff* = *max*{*h*(*A,B*),*h*(*B,A*)} and *h*(*A,B*) = *max*{ *min d*(*a, b*)} where A denotes the voxel set of the manually delineated volume, B denotes *a* ∈ *A b* ∈ *B* the voxel set of the predicted volume, and *d*(*a*, *b*) is the Euclidian distance between a and b. The Hausdorff distance was only estimated in-plane to avoid confounds from non-uniformly sampled data. The maximal Hausdorff distance (i.e., worst matching) across slices for each subject was then used for comparison. Superior performance was indicated by higher Dice, Jaccard, PPV, and SEN, and lower Hausdorff values. We also reported the computation time on a Linux-based [Red Hat Enterprise Linux Server release 7.4 (Maipo)] computing system (Intel E5-2680 v3 processor, 2.50 GHz, 256-GB RAM) for each method. The computation times reported do not include pre-processing steps (i.e., signal normalization, image resampling, and bias correction). Paired t-tests were used for statistical comparisons between different algorithms, and two-sample t-tests were used to compare T2w RARE and T2*w EPI images in each algorithm. The threshold for significance was set to the alpha level (*p* < 0.05).

## Results

Figure 2 illustrates the performance of our trained 3D U-Net64 algorithm compared to 2D U-Net64, 3D U-Net16, RATS, PCNN, and SHERM for rat brain segmentation in the CAMRI dataset. Across all measures except PPV (which had no significant differences), 3D U-Net64 showed superior T2w RARE brain segmentation performance over other existing methods (RATS, PCNN, and SHERM). Notably, although all the U-Net-based approaches produced ideal results with Dice > 0.90, the 3D U-Net64 showed significantly higher accuracy (p<0.05) versus the remaining U-Net approaches (2D U-Net64 and 3D U-Net16). The high PPV (>0.90) and low SEN (<0.90) from other existing methods (RATS, PCNN, and SHERM) indicate that brain segmentation was underestimated. In contrast, the low PPV (0.91 on anisotropic T2w RARE and 0.87 on anisotropic T2*w EPI) and high SEN (>0.95) from 2D U-Net16 suggest an overestimation. The significantly lower Hausdorff distance in 3D U-Net64 (4.29 on T2w RARE and 4.71 on T2*w EPI) further supports its superior ability to match the ground-truth. Notably, all U-Net methods provide excellent (Dice > 0.95) or high (Dice > 0.90) accuracy on T2w RARE and T2*w EPI images. All methods showed significant lower Dice in T2*w EPI images as compared to T2w RARE images (*p*<0.05). Specifically, the 3D U-Net64 still reached outstanding T2*w EPI brain segmentation performance metrics (Dice, Jaccard, PPV, and SEN > 0.95), whereas RATS, PCNN, and SHERM had lower T2*w EPI brain segmentation metrics (Dice, Jaccard, and SEN < 0.90). The compromised performance in the T2*w EPI image compared with T2w RARE indicates that these methods are less effective at handling data with lower spatial resolution. Further, 3D U-Net64 showed high accuracy (Dice > 0.95) in a validation study (Figure S2). Together, these results suggest that 3D U-Net64 is a reliable and reproducible approach for rat brain segmentation.

**Figure 2.**
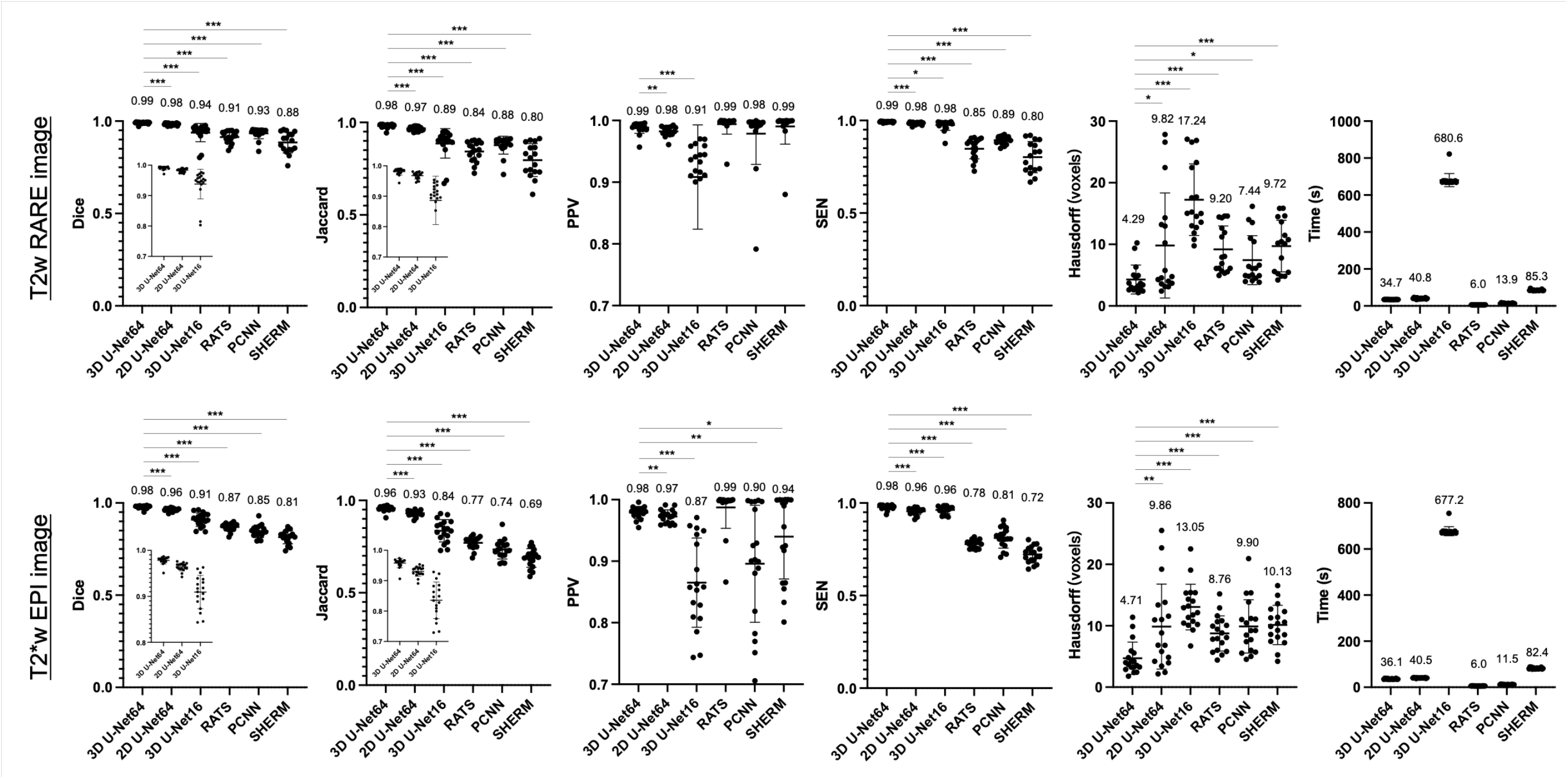
Brain segmentation performance metrics for 3D U-Net64, 2D U-Net64, 3D U-Net16, RATS, PCNN, and SHERM on the CAMRI T2w RARE (**upper row**) and T2*w EPI (**lower row**) data. Average value is shown above each bar. Two-tailed paired t-tests were used for statistical comparison between 3D U-Net64 with other methods (*p < 0.05, **p < 0.01, and *** p < 0.001).

Figures 3 and 4 illustrate the best and worst Dice score cases on T2w RARE images from the CAMRI dataset using all six methods. In the best case, both 2D U-Net64 and 3D U-Net16 provided nearly perfect segmentation (Dice > 0.98), and although the RATS, PCNN, and SHERM algorithms also showed high performance (Dice = 0.95), the inferior brain boundaries were less accurate. In the worst case, all U-Net methods still achieved a satisfactory segmentation with Dice > 0.90, but the RATS, PCNN, and SHERM algorithms failed to identify the brainstem, olfactory bulb, and inferior brain regions where the MRI signal was weaker.

**Figure 3.**
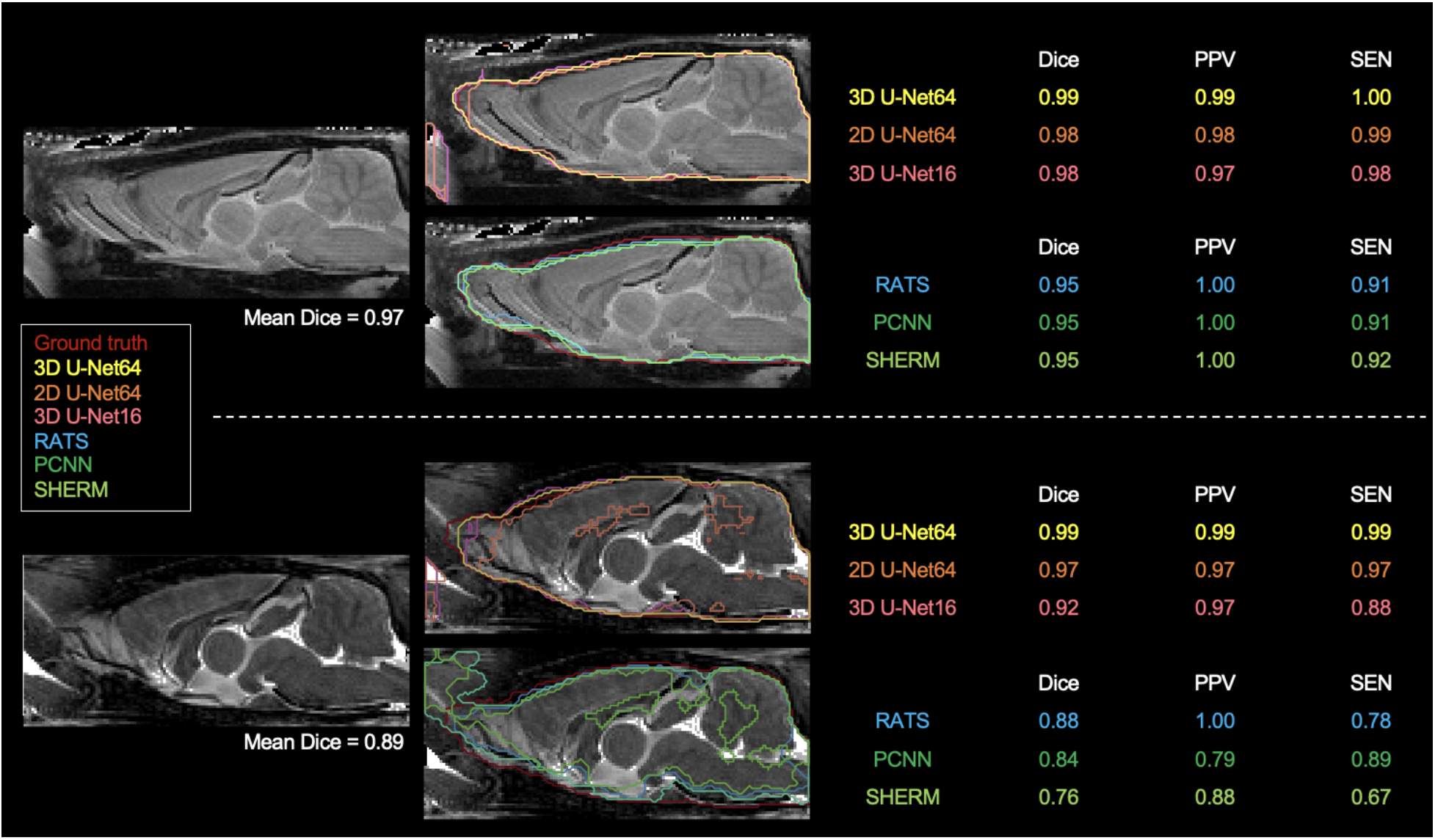
Best (upper panel) and worst (lower panel) segmentation comparisons for T2w RARE images. Selection was based on the highest and lowest mean Dice score (listed below the brain map) averaged over the six methods (3D U-Net64, 2D U-Net64, 3D U-Net16, RATS, PCNN, and SHERM). Anterior and inferior slices are more susceptible to error in RATS, PCNN, and SHERM, whereas all U-Net algorithms yield high similarly to the ground truth (all Dice > 0.90).

**Figure 4.**
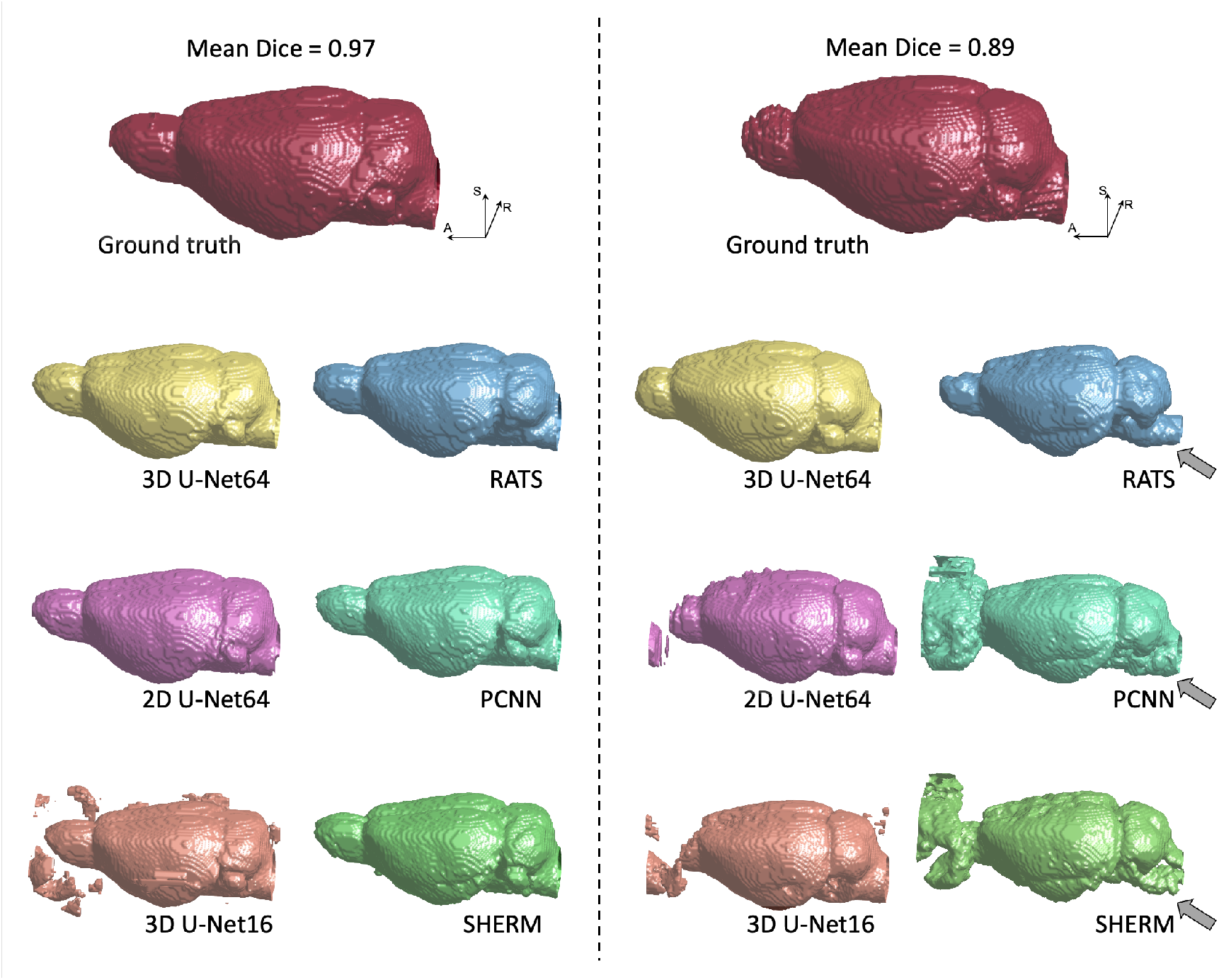
3D rendering of identified brain masks on the best and worst-case subjects for the T2w RARE rat dataset. Selection was based on the highest and lowest mean Dice score. Specifically, in the worst-case subject, 2D U-Net64, 3D U-Net16, and RATS missed the olfactory bulb, whereas PCNN and SHERM overestimated the olfactory bulb and incorporated surrounding frontal regions. Additionally, RATS, PCNN, and SHERM are missing significant portions of the cerebellum and brainstem (grey arrows). 3D U-Net64 and 3D U-Net32 produces excellent brain segmentation on both best and worst-case subjects.

Figure 5 illustrates the best and worst Dice score cases on T2*w EPI from the CAMRI dataset using all six methods. In the best case, all U-Net methods provide nearly perfect segmentation (Dice > 0.98). The RATS (Dice = 0.89), PCNN (Dice = 0.93), and SHERM (Dice = 0.83) algorithms showed acceptable segmentation performance but underestimated inferior brain regions. In the worst case, all U-Net methods still achieve a satisfactory segmentation with Dice > 0.90, but the RATS, PCNN, and SHERM algorithms failed to accurately identify the inferior brain boundaries and incorporated excessive tissue outside the brain (Dice < 0.85).

**Figure 5.**
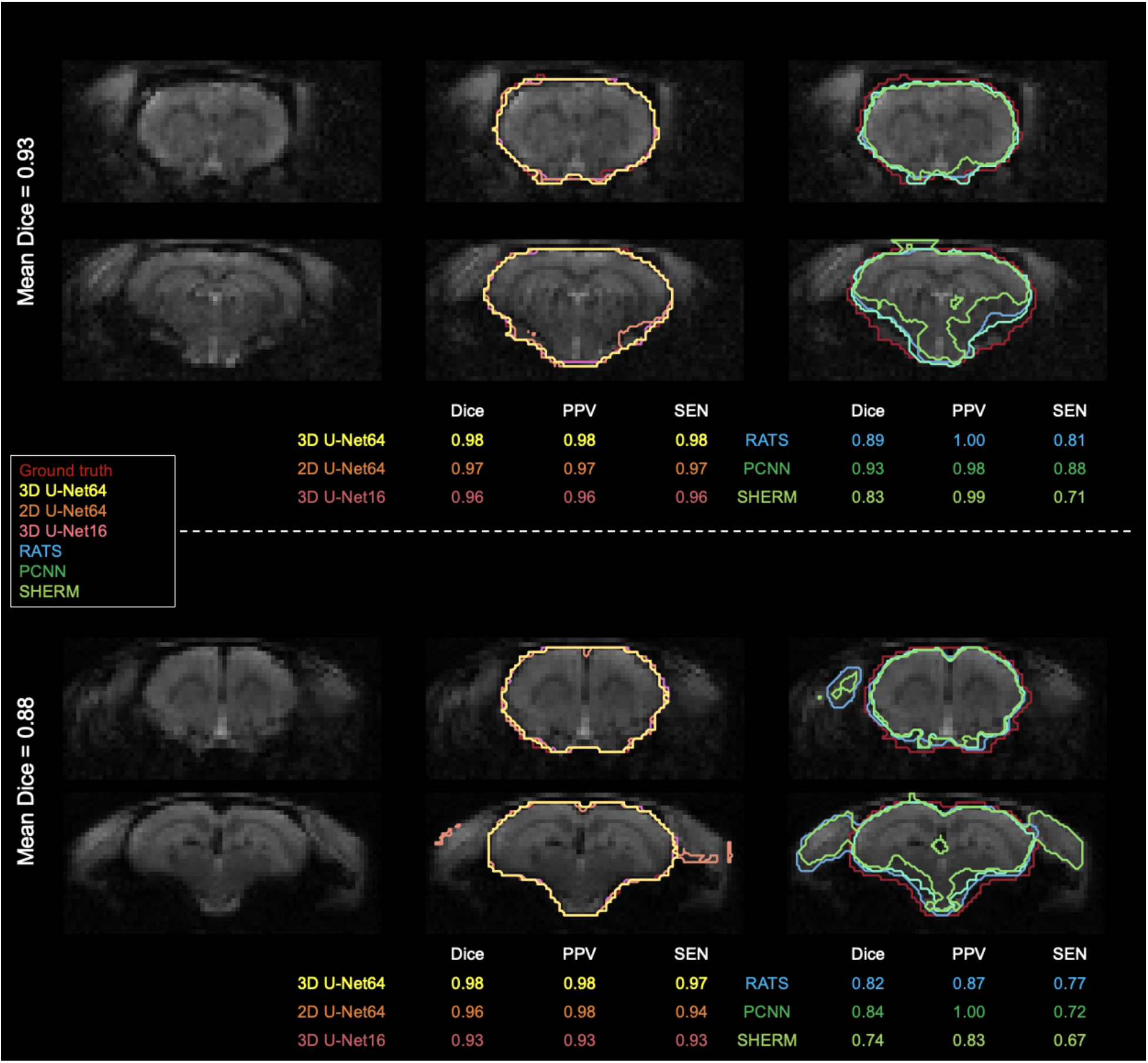
Best and worst segmentation comparisons for T2*w EPI images. Selection was based on the highest and lowest mean Dice score (listed above the brain map) averaged over the six methods (3D U-Net64, 2D U-Net64, 3D U-Net16, RATS, PCNN, and SHERM). Posterior and inferior slices are more susceptible to error in RATS, PCNN, and SHERM, whereas all U-Net algorithms are more similar to the ground truth (all Dice > 0.90).

We performed two validation analyses to evaluate the reliability of 3D U-Net segmentation performance. First, we added Gaussian noise to each testing image to evaluate 3D U-Net64 brain segmentation under different noisy environments. Figure 6 shows that the SNR and 3D U-Net64 segmentation performance decrease with increasing Gaussian white noise variance. The original mean SNR is about 26 for both T2w RARE and T2*w EPI data. When the SNR > 10 (i.e., reduced to 38% of original SNR) in T2w RARE images and SNR > 14 (i.e., reduced to 54% of original SNR) in T2*w EPI images, the segmentation accuracy could still reach an excellent Dice > 0.95. Figures S3 and S4 illustrate the best and worst examples on original T2w RARE and T2*w EPI images.

**Figure 6.**
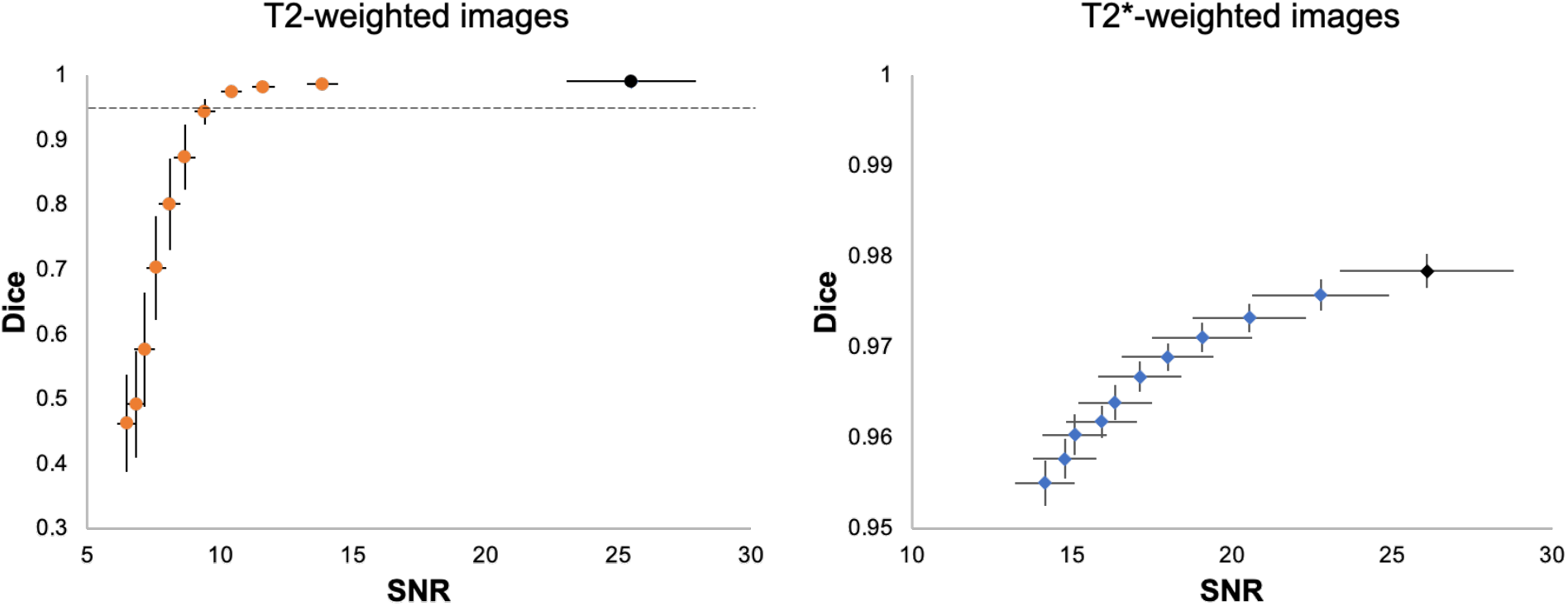
Segmentation performance of 3D U-Net64 with different image SNR. For each T2w (left) and T2*w image (right), we added noise with random Gaussian distribution in the normalized testing images with variance from 5×10^−5^ to 5×10^−4^ and increments of 5×10^−5^ to investigate the segmentation performance of 3D U-Net64. Black dots indicate the averaged SNR and Dice from the original images without adding noise. The horizontal dot line in (A) indicates a Dice of 0.95. Error bar represents the standard error of Dice and SNR.

Next, we evaluated the performance of 3D U-Net64 with regards to the model-training and model-validating sample sizes used during the model training process. We re-trained the 3D U-Net64 model with randomly selected training subgroups. Compared to the “standard model” trained on 55 subjects and validated on 14 subjects, the model with training rats < 25 showed significantly decreased accuracy for T2w RARE images, and the model with training rats < 50 showed significantly decreased accuracy for T2*w EPI images (Figure 7). However, it is worth noting that the model still reached excellent and stable segmentation performance (Dice > 0.95) with training subjects > 10 for both T2w RARE and T2*w EPI MRI data. There were no significant differences in each training sample selection across varied validation sample sizes in T2w RARE and T2*w EPI image segmentation, suggesting that a large number of model-validation subjects is perhaps unnecessary.

**Figure 7.**
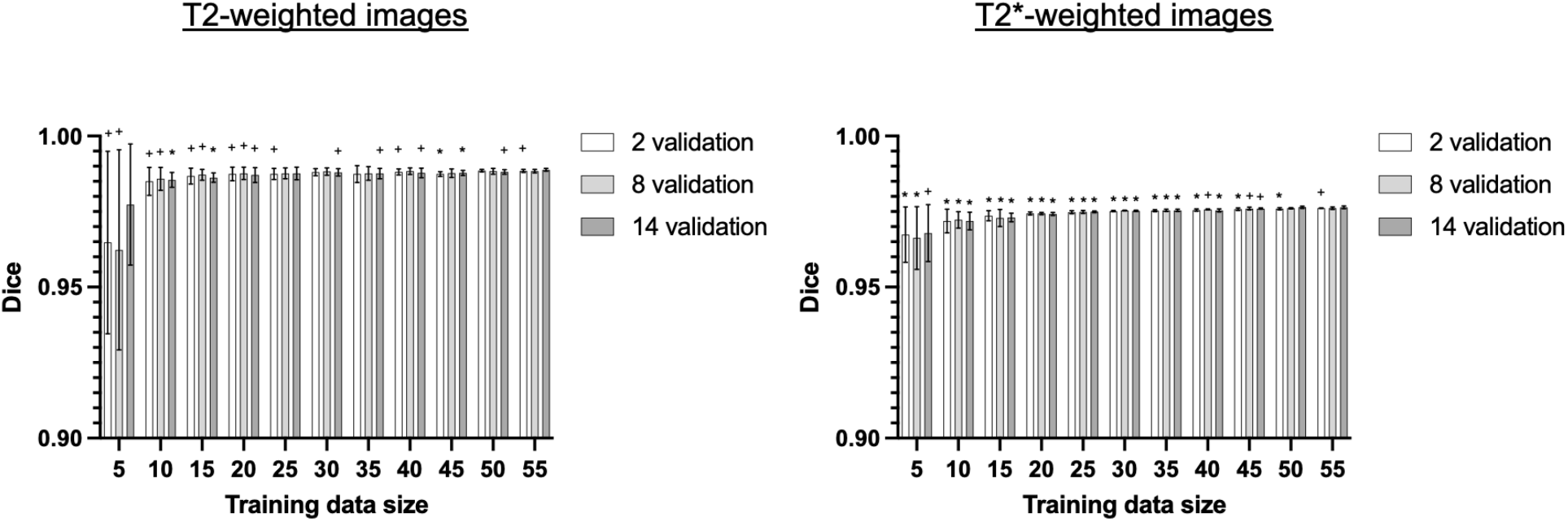
Segmentation performance of 3D U-Net64 across different model-training and model-validating sample sizes. The 3D U-Net64 was trained from randomly selected subgroups. In the training process, we randomly selected 5–55 training subjects in increments of 5 subjects from the total 55 training dataset, and 2, 8, and 14 validation subjects from the total 14 validation dataset. The random selection was repeated 5 times to avoid bias. Statistical analyses compared Dice values under various conditions against the Dice values obtained from 55 training rats and 14 validation rats (one tailed paired t-test, * p-< 0.05, + 0.05<p<0.1). No significant differences were found between various model-validation sample selections within each model-training data selection for both T2w RARE and T2*w EPI data (repeated measurement ANOVA).

## Discussion

U-Net is a neural network that is mainly used for classification and localization (Çiçek et al., 2016b; Ronneberger et al., 2015). In this study, we proposed a 3D U-Net architecture for automatic rat brain segmentation. Our results indicate that our proposed brain extraction framework based on 3D U-Net64 represents a robust method for the accurate and automatic extraction of rat brain tissue from MR images.

In this study where isotropic T2w RARE and T2* EPI data were segmented, RATS, PCNN, and SHERM still performed quickly and accurately (Dice > 0.8). 3D U-Net64 showed superior performance (average Dice = 0.99) over these methods and also outperformed our previously proposed 2D U-Net64 (Hsu et al., 2020) (please also refer to (Hsu et al., 2020) for detailed discussion about RATS, PCNN, and SHERM). The 3D U-Net algorithm remains “parameter free” in the segmentation process, as all parameters are automatically learned from the data itself. It should be noted that the U-Net architecture generally requires longer processing times and needs a higher level of computational power for model-training. For single patch inference, 2D U-Net takes less time than 3D U-Net. However, the patch size of 3D U-Net64 is one more dimension than that of 2D U-Net64, so to obtain the whole volume prediction, 3D U-Net needs less inference time and less time overall.

3D U-Net is a variant of 2D U-Net where the inputs are 3D volume (Çiçek et al., 2016a). It has the same encoder and decoder structure as in 2D U-Net. However, the encoder uses 3D convolution followed by 3D max-pooling to extract the features and the decoder uses 3D upsampling to reconstruct the annotated images (Çiçek et al., 2016a). The key advantage of 3D U-Net is its ability to utilize interslice contextual information. The 3D U-Net16 model tends to have higher SEN but lower PPV than the 2D U-Net64 model, as it misclassifies more background areas as brain (i.e., has more false positives). In other words, 2D U-Net64 may achieve higher precision because it makes fewer false positive errors. Although 3D U-Net16 can extract 3D context from the additional dimension that 2D U-Net64 lacks, its original 2D dimension resolution is also lost which means that the receptive field is decreased. Therefore, 2D U-Net64 achieves better performance than 3D U-Net16.

To illustrate the reliability of our proposed 3D U-Net64 architecture, we examined its accuracy under two conditions: conducting different Gaussian noise in the testing images and using a different number of rats from the original cohort in the training process. The results highlight the stability and robustness of 3D U-Net64 in segmenting the rat brain in both isotropic T2w RARE and T2*w EPI data. It is encouraging to observe excellent 3D U-Net64 performance (Dice > 0.95), even when the original image SNR is halved. Such performance is comparably higher than the conventional non-UNet-based methods using original SNR (RATS, PCNN, and SHERM). This demonstrates the noise-resistant capacity of 3D U-Net64 and suggests this approach may handle MRI data over a wide range of SNR quality. In addition, we also retrained the 3D U-Net64 model using a different number of subjects from the original cohort in the training process. This information is useful since the number of samples in a dataset are usually limited in rodent MRI studies. The Dice coefficient increases and stabilizes rapidly with > 10 training subjects for both T2w RARE and T2*w EPI. This finding demonstrates the utility of 3D U-Net64 with a limited number of training datasets.

There are several limitations of the 3D U-Net architecture. First, deep learning is a data driven classification, so segmentation accuracy highly relies on the training dataset. Because we trained our 3D U-Net algorithm by using only T2w RARE and T2*w EPI images in rats, additional training and optimization will be needed to segment brain images with different contrast (e.g., T1-weighted images). Second, deep learning methods require substantial amounts of manually labeled data (Verbraeken et al., 2020), and their performance can be affected by similarities between the training dataset and the data entering the U-Net brain segmentation process. While more training datasets of varying quality may further improve our current 3D U-Net model, our validation analysis suggested reliable segmentation performance can be reached with ≥ 10 training subjects. This indicates that the model can be established on a laboratory--or scanner-level basis. Third, our current 3D U-Net architecture uses a patch size of 64×64×64, which limits the testing image to an image matrix size of at least 64×64×64. Image resampling to a finer resolution is required if the image matrix size is smaller than 64×64×64. For a dataset at lower spatial resolution, a patch size of 16×16×16 as shown in this manuscript could be considered, although inferior performance may be expected. Alternatively, the source codes provided herein can be modified to adapt other patch sizes (such as 32×32×32) per the user’s discretion. For datasets with highly anisotropic resolution (e.g., high in-plane resolution with very few slices), we suggest that the users consider 2D U-Net as described in (Hsu et al., 2020).

## Conclusion

We proposed a 3D U-Net model, an end-to-end deep learning-based segmentation algorithm for brain extraction of 3D high resolution volumetric brain MRI data. The method is fully automated and demonstrates accurate brain mask delineations for isotropic structural (T2w RARE) and functional (T2*w EPI) MRI data of the rat brain. The method was compared against other techniques commonly used for rodent brain extraction as well as 2D U-Net. 3D U-Net shows superior performance in qualitative metrics including Dice, Jaccard, PPV, SEN, and Hausdorff distance. We believe this tool will be useful to avoid parameter-selection bias and streamline pre-processing steps when analyzing high resolution 3D rat brain MRI data. The 3D U-Net brain extraction tool can be found at https://camri.org/dissemination/software/.

## Data Availability Statement

The CAMRI rats dataset is available at https://doi.org/10.18112/openneuro.ds003646.v1.0.0. The 3D U-Net brain extraction tool can be found at https://camri.org/dissemination/software/.

## Acknowledgement

This work was supported by National Institute of Neurological Disorders and Stroke (R01NS091236, R44NS105500), National Institute of Mental Health (R01MH126518, RF1MH117053, R01MH111429, F32MH115439), National Institute on Alcohol Abuse and Alcoholism (P60AA011605, U01AA020023), and National Institute of Child Health and Human Development (P50HD103573). We thank Alicia Stevans at CAMRI for insightful discussion on this manuscript.

## Supporting information

**Figure S1.**
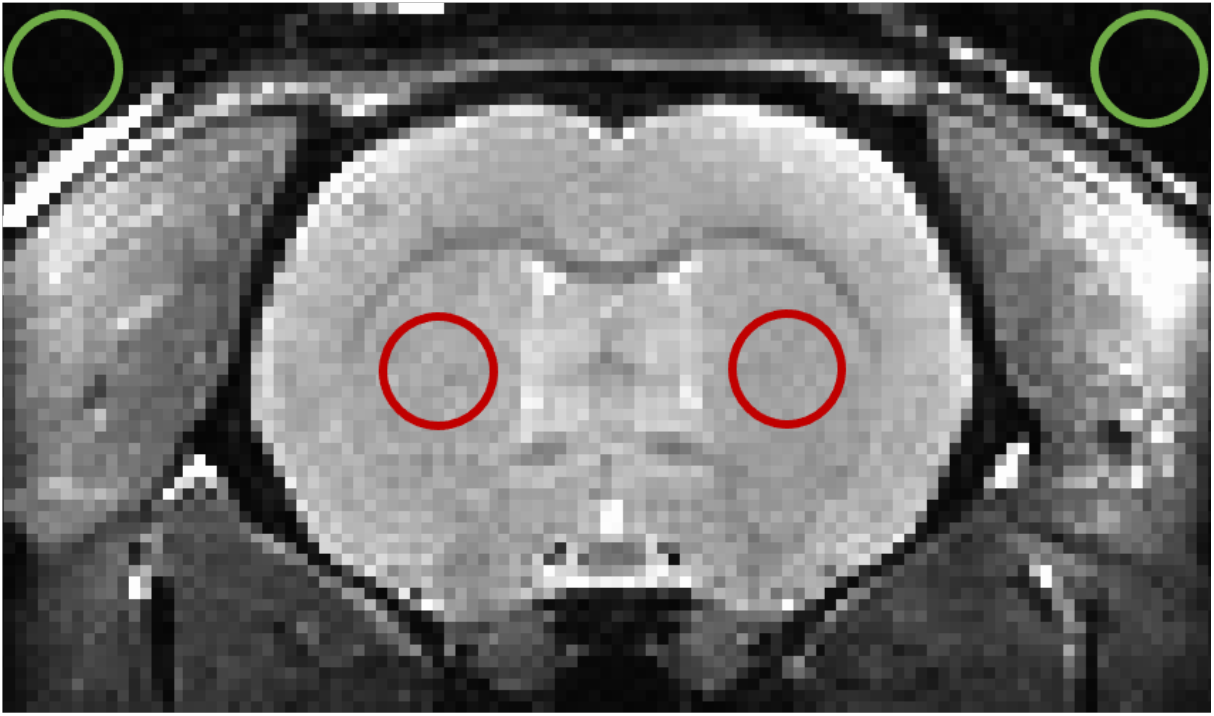
SNR calculation. The SNR was estimated to represent the image noise levels by calculating the ratio of the signal intensity in the area of interest (red circle) to that of the background (green circle).

**Figure S2.**
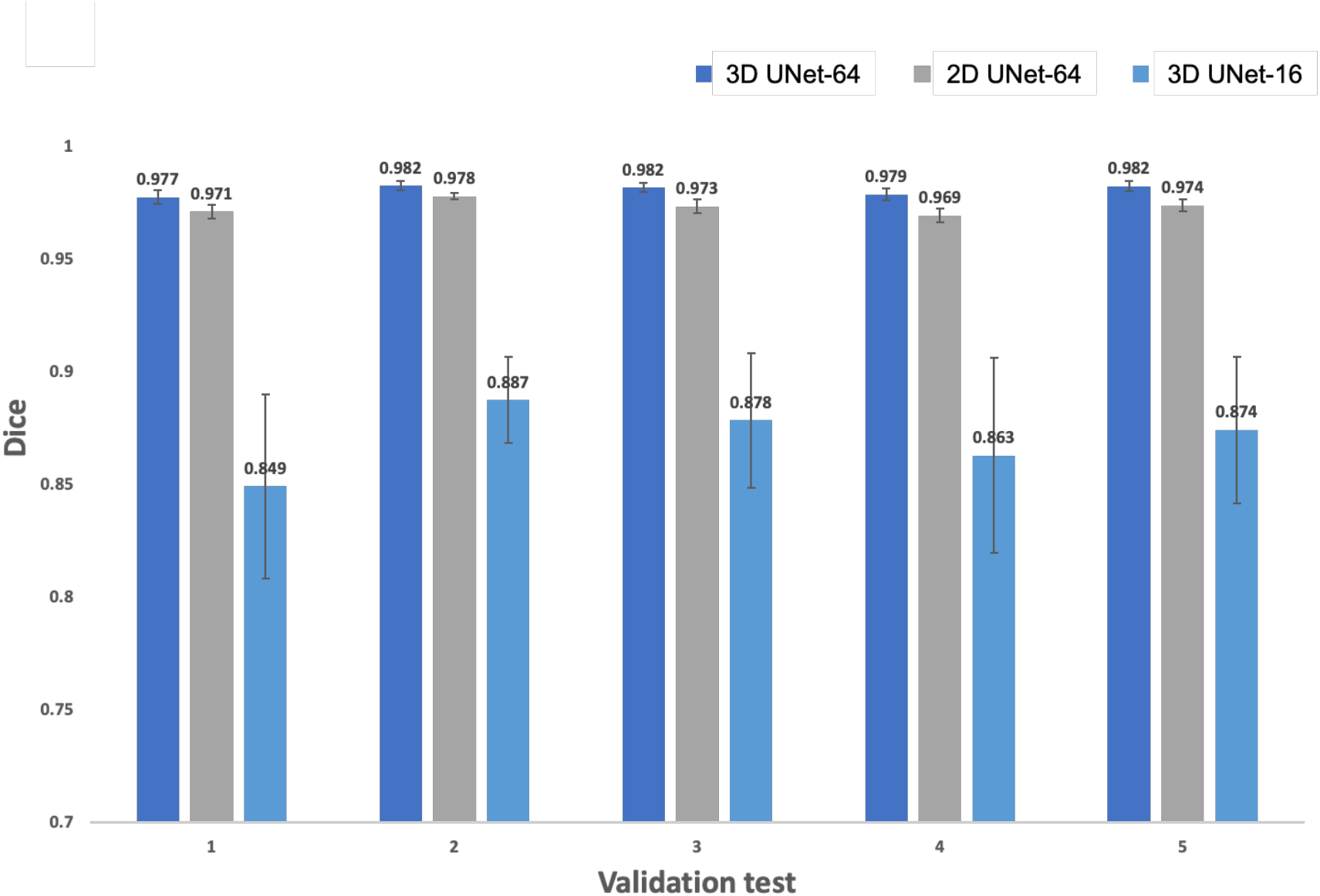
Performance validation results within the training process. In the training process, we randomly selected 80% of the rat data (55 rats) from the training dataset. The remaining 20% of the rat data (14 rats) from the training dataset was used for validating the U-Net model. We repeated this training-validation process five times to avoid randomness bias in data splitting. The U-Net model with highest averaged validation accuracy was then used as the final model for testing.

**Figure S3.**
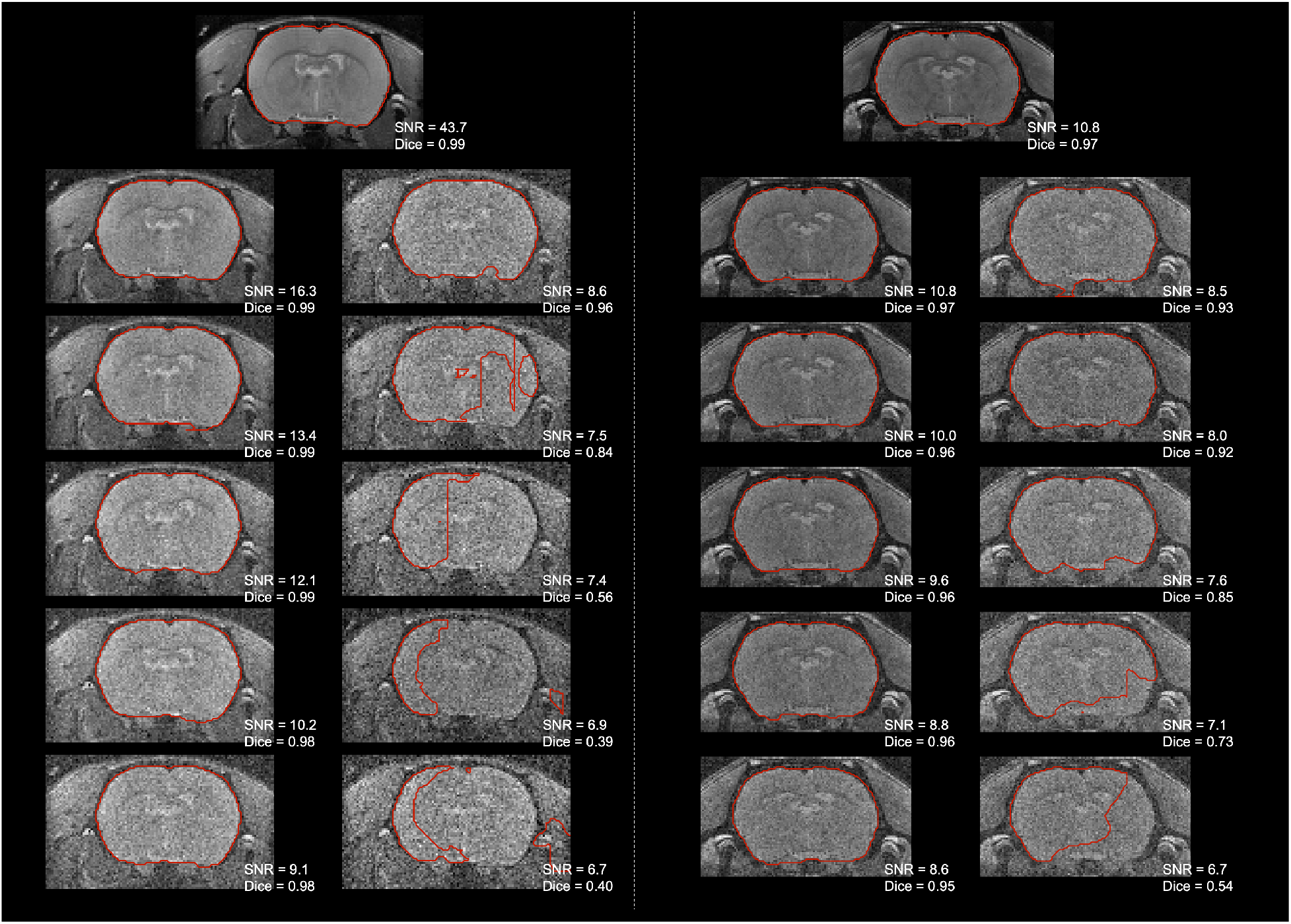
Highest (left panel) and lowest (right panel) Dice for T2w RARE images. For each rat, we added noise with random Gaussian distribution in the normalized testing images with variance from 5×10^−5^ to 5×10^−4^ in increments of 5×10^−5^. The SNR and segmentation accuracy (Dice) are labeled below each image.

**Figure S4.**
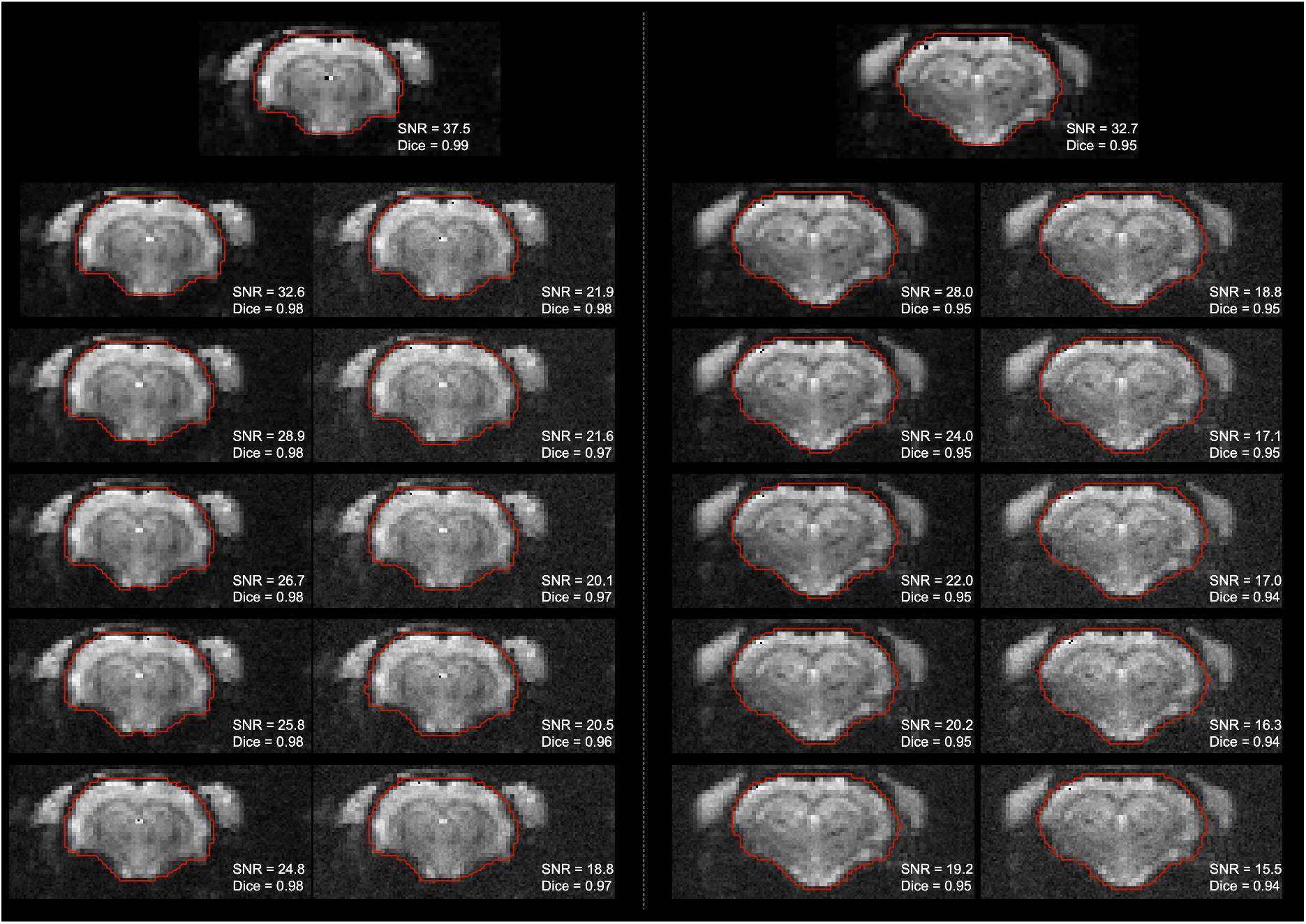
Highest (left panel) and lowest (right panel) Dice for T2*w EPI images. For each rat, we added noise with random Gaussian distribution in the normalized testing images with variance from 5×10^−5^ to 5×10^−4^ in increments of 5×10^−5^. The SNR and segmentation accuracy (Dice) are labeled below each image.

